# Embryonic motor neuron programming factors reactivate immature gene expression and suppress ALS pathologies in postnatal motor neurons

**DOI:** 10.1101/2024.04.03.587963

**Authors:** Emily R. Lowry, Tulsi Patel, Jonathon A. Costa, Elizabeth Chang, Shahroz Tariq, Hranush Melikyan, Ian M. Davis, Siaresh Aziz, Georgia Dermentzaki, Francesco Lotti, Hynek Wichterle

## Abstract

Aging is a major risk factor in amyotrophic lateral sclerosis (ALS) and other adult-onset neurodegenerative disorders. Whereas young neurons are capable of buffering disease-causing stresses, mature neurons lose this ability and degenerate over time. We hypothesized that the resilience of young motor neurons could be restored by re-expression of the embryonic motor neuron selector transcription factors ISL1 and LHX3. We found that viral re-expression of ISL1 and LHX3 reactivates aspects of the youthful gene expression program in mature motor neurons and alleviates key disease-relevant phenotypes in the SOD1^G93A^ mouse model of ALS. Our results suggest that redeployment of lineage-specific neuronal selector transcription factors can be an effective strategy to attenuate age-dependent phenotypes in neurodegenerative disease.

## Main Text

ALS is characterized by progressive motor neuron degeneration, typically leading to paralysis and death 2-5 years after diagnosis ^1^. Neurodegeneration in ALS is both cell type-specific and age-dependent, affecting only certain subtypes of motor neurons with a typical age of onset of 55-75 years. We have previously demonstrated that the expression of ∼7,000 genes and the accessibility of ∼100,000 chromatin regions change significantly over the course of postnatal motor neuron maturation ^2^. Considering the scale of these changes, it is challenging to pinpoint which genes or gene groups are responsible for the increased susceptibility of mature neurons to degeneration in ALS. We reasoned that reactivation of the transcriptional master regulators that control the immature motor neuron gene expression program might be a viable strategy to prevent or delay the onset of the disease.

We have previously shown that the majority of the genes induced in nascent motor neurons are direct targets of two transcriptional activators: ISL1 and LHX3 ^3–5^. ISL1 and LHX3 function as motor neuron selector transcription factors and can reprogram neural progenitors, pluripotent stem cells, and adult skin fibroblasts into immature spinal motor neurons ^4,6,7^. While essential for spinal motor neuron specification, both factors are downregulated in postnatal motor neurons (Fig. S1, A and B), and the LIM homeodomain motif that they bind becomes less accessible during postnatal motor neuron maturation ^2^. Therefore, we hypothesized that re-expression of these factors in postnatal motor neurons in vivo could re-activate an immature gene expression state, increase the ability of motor neurons to buffer intracellular stress, and reset motor neuron resistance to the deleterious effects of ALS-causing mutations.

## Results

### Chat enhancer for spinal motor neuron-specific re-expression of ISL1 and LHX3

To study the effects of postnatal ISL1 and LHX3 reactivation in skeletal motor neurons, we generated adeno-associated viruses (AAVs) that drive expression of *Isl1* and *Lhx3* under the control of ChatE, a 1000 bp enhancer located 3 kb upstream of the *Chat* gene encoding choline acetyltransferase (Fig. 1A). We identified ChatE in a temporal ATAC-seq dataset ^2^ as a chromatin region that is continuously accessible in spinal motor neurons throughout postnatal life (Fig. 1A) and contains binding sites for a suite of transcription factors that control motor neuron specification and maturation (Fig. S1C) ^2,4,5^. To evaluate the efficiency and specificity of ChatE-driven expression, AAV-Isl1 and AAV-Lhx3 were co-administered at a 1:1 ratio by intracerebroventricular injection in neonatal mice at postnatal day 1 (P1; Fig. 1B). Strong, motor neuron-specific expression of ISL1 and LHX3 was observed by immunohistochemistry (IHC) in the lumbar spinal cord within 7 days post-injection (Fig. 1C and Fig S1, B and D). Other local CHAT-expressing cells, such as V0c interneurons, remained ISL1 and LHX3 negative (Fig. 1C). Under both low titer (6-9E+10 vg/animal) and high titer transduction conditions (3-4E+11 vg/animal), we observed that ∼90% of CHAT+ motor neurons in the lumbar spinal cord were AAV+ at early post-injection time points (7-14 days), with near-complete overlap between ISL1- and LHX3-positive cells (Fig. 1C). Under low titer conditions, the percentage of AAV-expressing motor neurons dropped to ∼22% by P45, with 11% co-expressing ISL1 and LHX3, 2% expressing ISL1 alone, and 9% expressing LHX3 alone (Fig. 1D). Under high titer conditions, LHX3 expression appeared to stabilize, while ISL1 continued to decrease with time (Fig. S1D). By P45, 84% of motor neurons retained expression of AAV-driven LHX3, with 23% co-expressing ISL1 and LHX3 and 61% expressing LHX3 alone (Fig. 1D). AAV-Isl1+AAV-Lhx3 treatment appeared to be well-tolerated across all titers and timepoints, and did not result in any overt behavioral changes, motor or otherwise, compared to AAV-mCherry-treated or untreated controls.

**Fig. 1:**
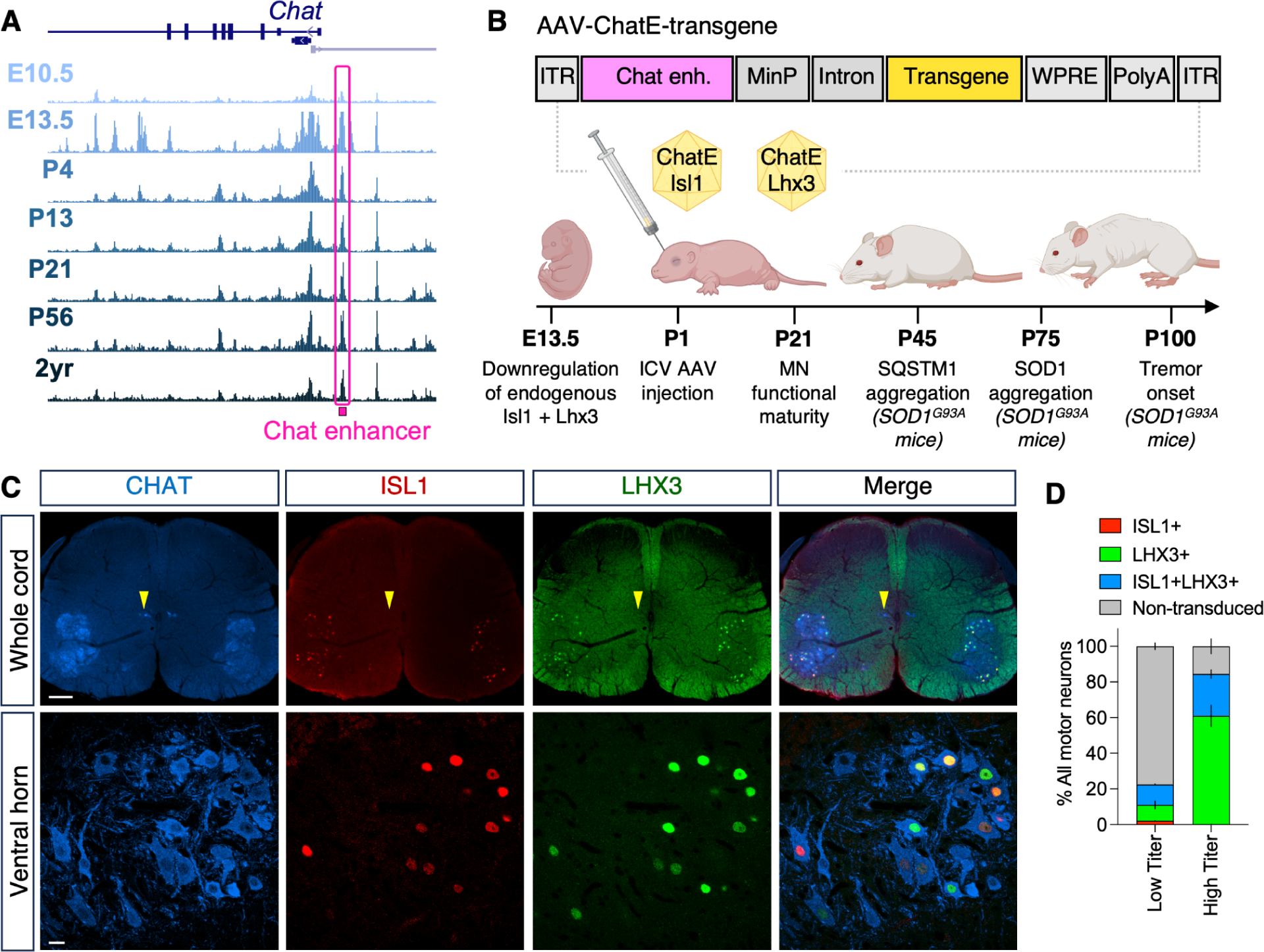
Design and in vivo validation of ChatE-driven Isl1 and Lhx3 AAVs. (A) Schematic of ATAC-seq data from ^2^ showing accessibility of ChatE (pink) in spinal motor neurons over time. (B) Design of AAV-ChatE constructs and experiments for evaluating them in vivo. (C) Immunofluorescent staining of ectopic ISL1 and LHX3 in the L4-L5 region of the spinal cord in a P14 mouse. Ectopic ChatE-driven ISL1 and LHX3 expression is restricted to CHAT+ motor neurons in the ventral horn (first row, 5x epifluorescence; scale bar represents 100 µm), and most transduced motor neurons are double-positive for ISL1 and LHX3 (second row, 40x confocal; scale bar represents 10 µm). Arrowheads indicate cholinergic V0c interneurons, which remain negative for ISL1 and LHX3 (D) Distribution of ISL+, LHX3+, ISL+LHX3+, and non-transduced cells among CHAT+ cells in the L4-L5 ventral horn at P45 from animals treated with low titer (6-9E+10 vg/animal) or high titer AAVs (3-4+11 vg/animal). Mean percentage of motor neurons in each condition was quantified from 6 hemisections per animal and 15 confocal images per hemisection from 2 (low titer) or 6 (high titer) animals. Error bars represent SEM.

### Global gene expression and chromatin accessibility profiling of transduced neurons

To evaluate whether heterochronically re-expressed ISL1 and LHX3 could reactivate their embryonic targets, we stained lumbar spinal cord tissue collected at P45 for MNX1 (HB9), a gene that is specifically expressed in nascent spinal motor neurons and is downregulated before birth (Fig. S2A). We found that MNX1 was virtually undetectable in untreated or control AAV-treated animals (Fig. 2A). Treatment with AAV-Isl1+AAV-Lhx3 led to significant upregulation of MNX1 (p<0.0001; Fig. 2, A and B), with 50% AAV-Isl1^+^ motor neurons and 24% of motor neurons overall exhibiting MNX1 expression.

**Fig. 2:**
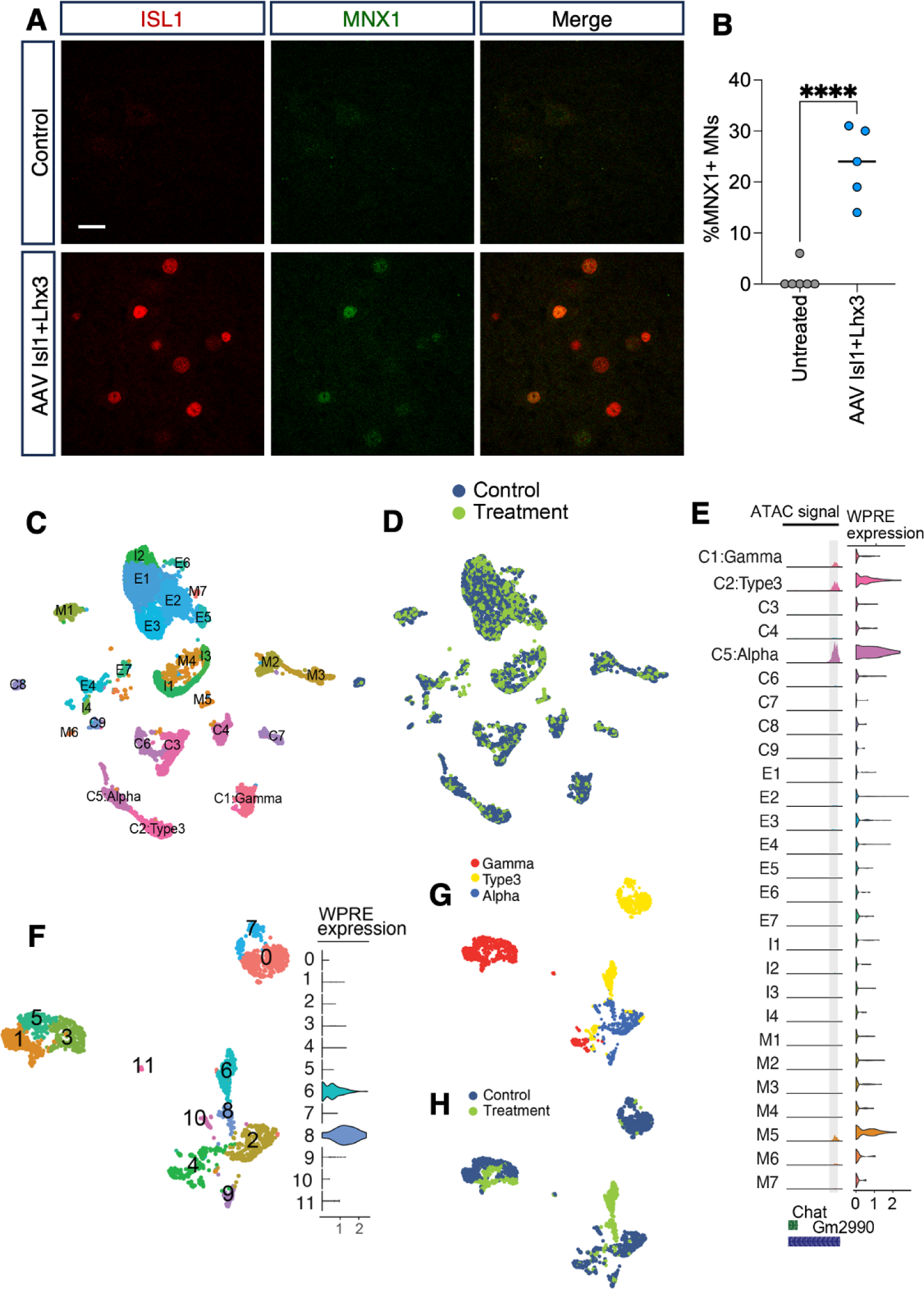
ISL1 and LHX3 re-expression drive motor neuron subtype-specific changes in gene expression. (A) Immunostaining for the ISL1+LHX3 target MNX1 in spinal motor neurons in the ventral horn in P45 animals. MNX1 is undetectable in untreated animals but is upregulated in ISL+ motor neurons in animals treated with AAV-Isl1+AAV-Lhx3. Images are maximum intensity projections of 15 z sections taken 3 µ apart. Scale bar represents 10 µm. (B) Quantification of MNX1+ motor neurons in the L4-L5 region of the spinal cord in untreated or AAV-Isl1+AAV-Lhx3 animals at P45. The percentage of MNX1+ cells among all CHAT+ cells was quantified in 15 confocal images from 1 (untreated) or 6 (AAV-treated) 70 µ hemisections per animal. Each point represents percentage per hemisection (untreated) or mean percentage per hemisection (AAV-treated) from one animal. Black bars represent animal medians per treatment condition. Significance determined by unpaired two-tailed t test (p=0.0004, t=5.164, df=10). (C) Clustering of single nuclei multiome dataset using Canonical Correlation Analysis (CCA)-based integration of snRNA-seq data. Clusters are labeled based on expression of Chat (C=cholinergic), Slc17ac6 (E=excitatory), Gad1 (I =inhibitory), or >1 of these three genes (M=multiple). The three motor neurons clusters are labeled Alpha, Gamma, or Type 3 based on known subtype markers. (D) Each cluster is populated by both control and treatment cells. (E) Left: Normalized ATAC signal (range 0 - 4700) at the ChatE in all clusters. High signal is apparent in C1:Gamma motor neurons, C2: Type 3 motor neurons, C5: Alpha motor neurons and cluster M5. Right: Violin plots showing expression of WPRE transcripts from the AAV. The highest levels of expression are seen in alpha and type 3 motor neurons and in cluster M5. (F) Re-clustering of C1: Gamma, C2: Type 3, and C5: Alpha motor neurons clusters from (A) without the use of integration anchors. WPRE expression is highest in clusters 6 and 8 (G-H) Treatment gamma cells are found in the same clusters with control gamma cells. However, most of the treatment alpha and type3 cells form independent clusters (alpha prime and type 3 prime, respectively), separate from clusters populated largely by control alpha and type 3 cells.

The reactivation of MNX1 prompted us to evaluate the global effects of ISL1 and LHX3 re-expression on postnatal motor neuron gene expression and chromatin accessibility by performing single nucleus multiome RNA and ATAC sequencing in the spinal cord. To enrich for motor neuron nuclei in whole spinal cord samples, we used Chat-Cre;Sun1-sfGFP-Myc (INTACT) mice that express a nuclear envelope-bound GFP reporter driven by *Chat* ^2,8^. INTACT mice were either injected at P1 with AAV-Isl1+AAV-Lhx3 (Treatment group; n=8 mice) or left untreated (Control group; n=6 mice). GFP+ nuclei from both conditions were then isolated and sequenced at P21. RNAseq reads were aligned to a custom genome that included four additional elements present in the AAV constructs to identify AAV+ nuclei within the treatment group: the human ISL1 and LHX3 transgene-coding sequences, the chimeric intron, and the WPRE 3’UTR element (Fig. 1B). Canonical correlation analysis (CCA)-based integration of the two datasets resulted in the identification of 27 clusters (Fig. 2C) populated by nuclei from both control and experimental animals (Fig. 2D and Fig. S2B). Using motor neuron markers defined by previous single nuclei RNAseq studies ^8,9^, we identified three clusters corresponding to alpha motor neurons (C5), gamma motor neurons (C1), and type 3 motor neurons (C2) (Fig. 2C and Fig. S2, C and D). Consistent with the motor neuron specificity of ChatE, expression of AAV transgenes was largely restricted to the motor neuron clusters (Fig. 2E): 90.5% of alpha motor neuron nuclei, 70.9% of type 3 motor neuron nuclei, and 26.5% of gamma motor neuron nuclei from the treatment group expressed at least one of the four AAV elements (Fig. 2E). We identified one additional non-motor neuron cluster (M5) with 55% of nuclei expressing AAV transcripts. ATACseq analysis confirmed that the three motor neuron clusters, as well as the M5 cluster, exhibited high levels of accessibility at the ChatE locus (Fig. 2E).

To understand the impact of treatment on gene expression, we performed differential gene expression analysis for each of the four AAV-expressing clusters. Within each cluster, all AAV-expressing nuclei were compared with an equal number of randomly selected nuclei from the control group. This analysis showed that >200 genes were differentially expressed in both alpha and type 3 clusters, while fewer than 20 genes were differentially expressed in the gamma and M5 clusters (Fig. S2E). The absence of gene expression changes in gamma motor neurons was especially surprising, given that ISL1 and LHX3 proteins can be detected in these cells by IHC (Fig. S3). Together, these data suggest that the effects of heterochronically re-expressed ISL1 and LHX3 are not only specific to motor neurons over other cell types but are also selective among motor neuron subtypes.

### Motor neuron subtype-specific effects of ISL1 and LHX3 on gene expression

Considering the broad impact of AAV-Isl1+AAV-Lhx3 on gene expression on alpha and type 3 motor neurons, we asked whether selectively re-clustering motor neuron nuclei without CCA-based integration would result in the segregation of control vs. treatment nuclei (Fig. 2F-H). Indeed, we found that most type 3 and alpha motor neuron nuclei in the treatment group formed unique clusters (6 and 8, respectively) that separated from the type 3 (0 and 7) and alpha (2, 9, and 10) motor neuron clusters populated primarily by control nuclei (Fig. 2, G and H, and Fig. S4A). We termed these unique clusters alpha prime and type 3 prime. In contrast, gamma motor neuron nuclei from both conditions populated the same clusters (Fig. 2, G, and H), consistent with the minimal gene expression changes observed in these cells (Fig. S2E). To ensure that the separation of alpha prime and type 3 prime clusters from their respective control clusters was due to biological rather than technical reasons, we used the same methods to re-cluster the non-motor neuron cholinergic nuclei and found that all resulting clusters had similar proportions of control and treatment nuclei (Fig. S4, B, C, and D).

The formation of treatment-specific clusters allowed us to focus our subsequent analyses on the cells most impacted by ISL1 and LHX3 re-expression. Differential gene expression analysis identified 86 genes significantly upregulated and 119 genes significantly downregulated in alpha prime vs. the remaining alpha nuclei (referred to hereafter as alpha differentially-expressed genes (DEGS); Fig. 3A), and 253 genes significantly upregulated and 117 genes significantly downregulated in type 3 prime vs the remaining type 3 nuclei (type 3 differentially-expressed DEGs; Fig. 3A). Only 88 genes, or 15% of total DEGs, were shared between the alpha and type 3 motor neurons, indicating that the effects of transgene expression were largely subtype-specific even among the cell populations that are strongly AAV-positive. To independently validate the single nuclei differential gene expression, we performed bulk RNA-seq analysis of GFP+ cells isolated from P21 INTACT control mice and mice treated with AAV-Isl1+AAV-Lhx3 at P1. Despite the lower sensitivity of the bulk sequencing dataset, we found a significant positive correlation between the expression of both the alpha and type 3 DEGs in the single nuclei and bulk sequencing datasets (alpha DEG R^2^=0.40, p<0.0001; type 3 DEG R^2^=0.21, p<0.0001; Fig. S5). Finally, we used IHC to confirm the differential expression of CADPS2, one of the top induced genes in alpha prime neurons. Staining of lumbar spinal cord sections from an independent set of control and AAV-Isl1+AAV-Lhx3 treated mice revealed a significant upregulation and striking perinuclear accumulation of CADPS2 protein in transduced ISL1+ motor neurons (Fig 3, B and C).

**Fig. 3:**
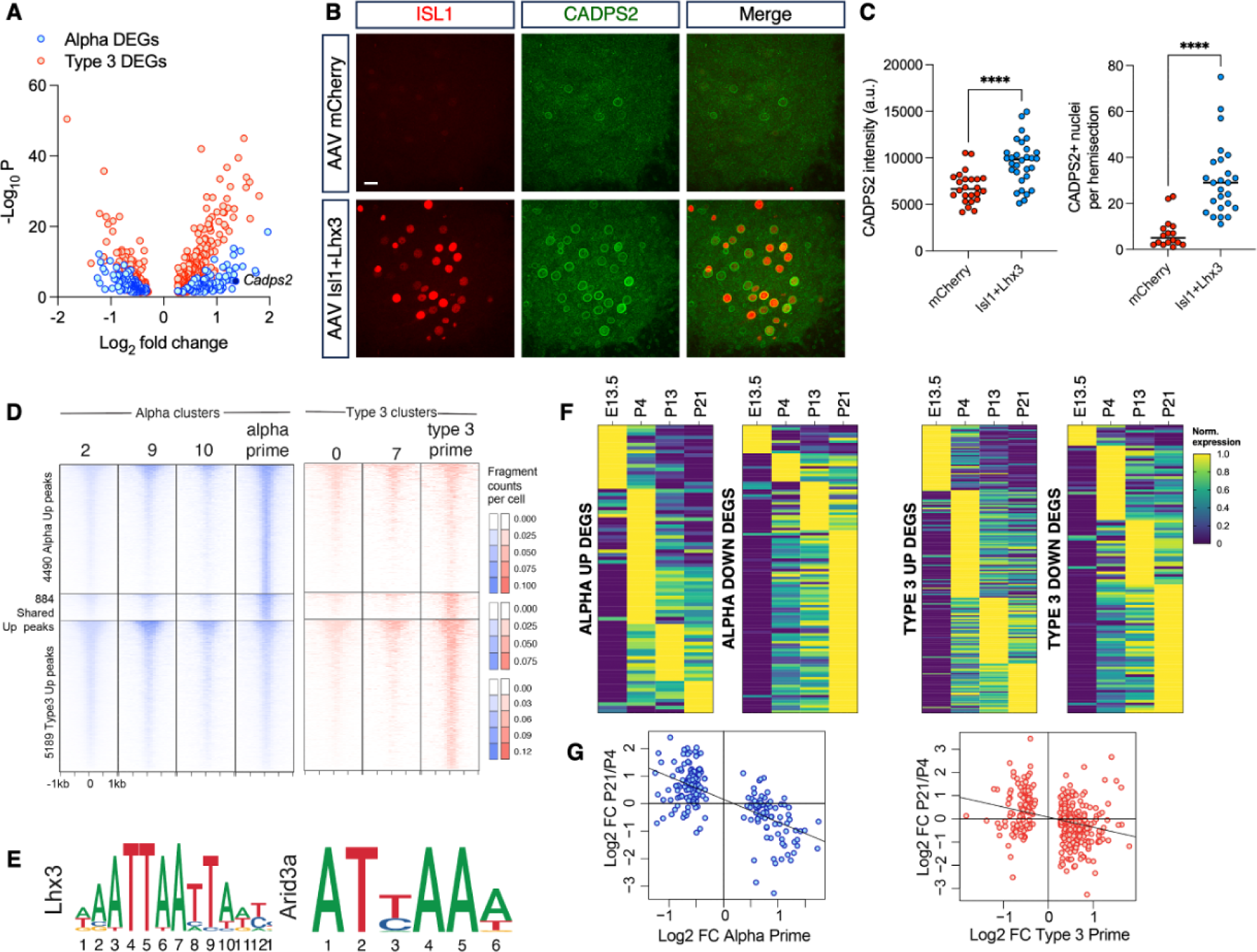
Gene expression changes in unique alpha and type 3 clusters are consistent with more youthful state. (A) Plot of DEGs with log2 fold-change of at least 0.25 and an adjusted p value <0.05 for alpha motor neuron-specific cluster 8 compared to clusters 2, 9, and 10 (blue), and for type 3 motor neuron-specific cluster 6 compared to clusters 0 and 7 (pink). Each point represents one gene. The alpha motor neuron-specific DEG Cadps2 is indicated in dark blue. (B) Representative immunostaining for ISL1 (red) and CADPS2 (green) in AAV-mCherry and AAV-Isl1+Lhx3-treated animals. AAV-Isl1+AAV-Lhx3 treatment leads to up-regulation of CADPS2 in AAV+ cells. Images are maximum intensity projections of 15 z sections taken 3 µ apart. Scale bar represents 10 µm. (C) Quantification of CADPS2+ nuclei per hemisection and overall CADPS2 signal intensity. Each point represents data from one hemisection, with 6 hemisections assessed per animal, n=4 (AAV-mCherry) or n=5 (AAV-Isl1+Lhx3) animals per treatment condition. Black bars represent median values. Significance determined by unpaired two-tailed t test (CADPS2+ nuclei p<0.0001, t=5.479, df=39; CADPS2 intensity p<0.0001, t=4.532, df=52). (D) Heatmaps of ATAC-seq reads at genomic regions that gain accessibility only in treatment alpha clusters (top), only in treatment type 3 clusters (bottom) or in both treatment alpha and type 3 clusters (middle). (E) The top 2 motifs among upregulated peaks are the LHX3 binding motif and a related homeodomain motif. (F) Alpha and type 3 DEGs from (A) plotted to show the timepoint at which they reach maximal expression during normal development based the longitudinal bulk sequencing dataset from ^2^. Expression values for each gene are normalized so that minimum value between E13.5 and P21 is represented as 0, and maximum value is 1. (G) Correlations for the alpha and type 3 DEGs between AAV-Isl1-Lhx3-mediated changes in gene expression (from (A)) vs. maturation-mediated changes in expression between P21 and P4 (from ^2^), suggesting that alpha prime and type 3 prime motor neurons are reverted to an early postnatal state. For alpha DEGs, R2=0.4963, p<0.0001. For type 3 DEGs, R2=0.07469, p<0.0001.

### Putative regulatory elements that gain accessibility are enriched for LHX3-HD binding motif

Cooperative binding of ISL1 and LHX3 to DNA in differentiating embryonic stem cells results in a transient increase in chromatin accessibility that is rapidly lost following *Lhx3* downregulation in maturing motor neurons ^5^. To investigate whether re-expression of these factors in postnatal neurons changed chromatin accessibility at P21, we examined the single nucleus ATACseq dataset. We first performed differential peak analysis, which led to the identification of 5374 upregulated and 2981 downregulated peaks (among ∼89,000 total accessible sites) in alpha prime compared to alpha nuclei, and 6073 upregulated and 2644 downregulated peaks (among ∼87,000 total accessible sites) in type 3 prime compared to type 3 nuclei (Figure 3D, upregulated peaks shown). Next, we asked which transcription factor motifs were enriched in the regions that gained or lost chromatin accessibility following ISL1 and LHX3 re-expression. For both alpha prime and type 3 prime motor neurons, the top motifs enriched in upregulated peaks, but not downregulated peaks, were Lhx3 and Lhx3-like LIM homeodomain motifs (Fig. 3E). However, despite the enrichment of the LHX3 motif across both clusters, there was only minimal overlap (∼8%) of the chromatin domains that gained accessibility between each cluster, which echoed the subtype-specificity of the DEGs. These findings suggest that re-expressed ISL1 and LHX3 increase chromatin accessibility at genomic loci that harbor canonical LHX3 binding motifs, which parallels their activity during normal motor neuron development ^4^, but that their activity in postnatal motor neurons is subtype-specific.

To further explore the relationship between changes in chromatin accessibility with changes in gene expression, we analyzed the distance between the differential peaks and the DEGs for each motor neuron subtype. We found that 70% of upregulated alpha DEGs and 87% of upregulated type 3 DEGs were proximal to at least one chromatin region that gained accessibility, compared to only 32% and 22% of downregulated alpha and type 3 DEGs. Together, the observations that Lhx3 motifs are preferentially enriched in upregulated peaks, and that upregulated peaks are preferentially associated with upregulated genes, suggest that ISL1 and LHX3 act as transcriptional activators when they are re-expressed postnatally, much like they do during development ^3,5,6^.

### Activation of early postnatal gene expression program in transduced motor neurons

Encouraged by the major changes in gene expression and accessibility observed in AAV-transduced cells, we set out to evaluate whether heterochronic ISL1 and LHX3 re-expression was preferentially activating genes expressed in immature motor neurons. As a benchmark for transcriptional “age” of the alpha and type 3 prime nuclei, which were collected at P21 from animals treated at P1, we used longitudinal motor neuron transcription profiles generated by bulk sequencing of purified INTACT nuclei at E13.5, P4, P13, and P21 ^2^. We first examined the normal temporal expression patterns of all alpha and type 3 DEGs. Among both motor neuron subtypes, we found that most of the genes that were upregulated in response to AAV-Isl1+AAV-Lhx3 treatment normally peak during embryonic and early postnatal ages (E13.5 or P4; Fig. 3F). Conversely, genes that were downregulated in response to AAV-Isl1+AAV-Lhx3 treatment are normally most highly expressed as motor neurons approach functional maturity (P21; Fig. 3F). We then compared AAV-driven changes in the alpha and type 3 DEGs vs. maturation-driven changes in gene expression between P21 and E13.5, P4, or P13. We found significant negative correlations across all comparisons, with the strongest correlation among genes that are differentially expressed between P21 and P4 (Fig. 3G and Fig. S6A). Together, these data suggest that re-expressed ISL1 and LHX3 activate a gene expression subprogram that is characteristic of perinatal spinal motor neurons.

This idea was further underscored by gene ontology (GO) analyses of the upregulated alpha and type 3 DEGs, where the most significantly enriched terms for both cell types fell into categories relating to neural development, synaptogenesis, and axonogenesis (Fig. S6, B through D). Indeed, many of the shared, upregulated alpha and type 3 DEGs have well-established roles in motor neuron development, including *Sema3c* ^10^, *Epha3* ^11,12^ and *Robo1* ^13^. Thus, while ISL1 and LHX3 re-expression yielded a host of cell type-specific DEGs, it appeared to have the same global effect of promoting a less mature state on both alpha and type 3 motor neurons.

### ISL1 and LHX3 re-expression prevent the formation of SQSTM1-positive protein aggregates in a mouse model of ALS

To probe the impact of reactivating perinatal motor neuron genes on late-onset motor neuron degeneration, we evaluated AAV-Isl1 and AAV-Lhx3 in the SOD1^G93A^ transgenic mouse model of ALS, where most motor neuron pathologies occur in mature cells. This mouse model recapitulates many biochemical and behavioral features of the disease, including severe deficits in the processing of misfolded proteins in vulnerable spinal motor neuron populations, followed by neuromuscular junction denervation, motor neuron degeneration, neuroinflammation, and clinical symptoms that progress swiftly until endstage at ∼P157.

One of the earliest histological markers of disease in this model is dysregulation of SQSTM1, a key component of both the ubiquitin-proteasome system (UPS) and the macroautophagy pathway that regulates lysosomal protein degradation ^14^. Large, round aggregates of SQSTM1 (termed “round bodies” ^15^) are detectable in the cytoplasm of lumbar motor neurons by P35 ^16^, and by P45, they can be found in nearly one-third of motor neurons (see Fig. 4A). Highlighting the relationship between proteostatic defects and clinical severity, motor neuron-specific knockout of the autophagy gene Atg7 in SOD1^G93A^ mice enhances SQSTM1 aggregation and hastens the onset of clinical symptoms ^15^. Overexpression of SQSTM1 itself promotes the cytoplasmic aggregation of mutant SOD1 and accelerates the onset of clinical phenotypes in SOD1^H46R^ mice ^17^. Importantly, SQSTM1-positive aggregates that are co-positive for TDP43 and other ubiquitinated proteins are commonly observed in postmortem spinal cord tissues from both familial and sporadic ALS patients ^18^, and mutations in SQSTM1 have been causally linked to ALS ^19^, reinforcing the translational relevance of SQSTM1 dysregulation.

**Fig. 4:**
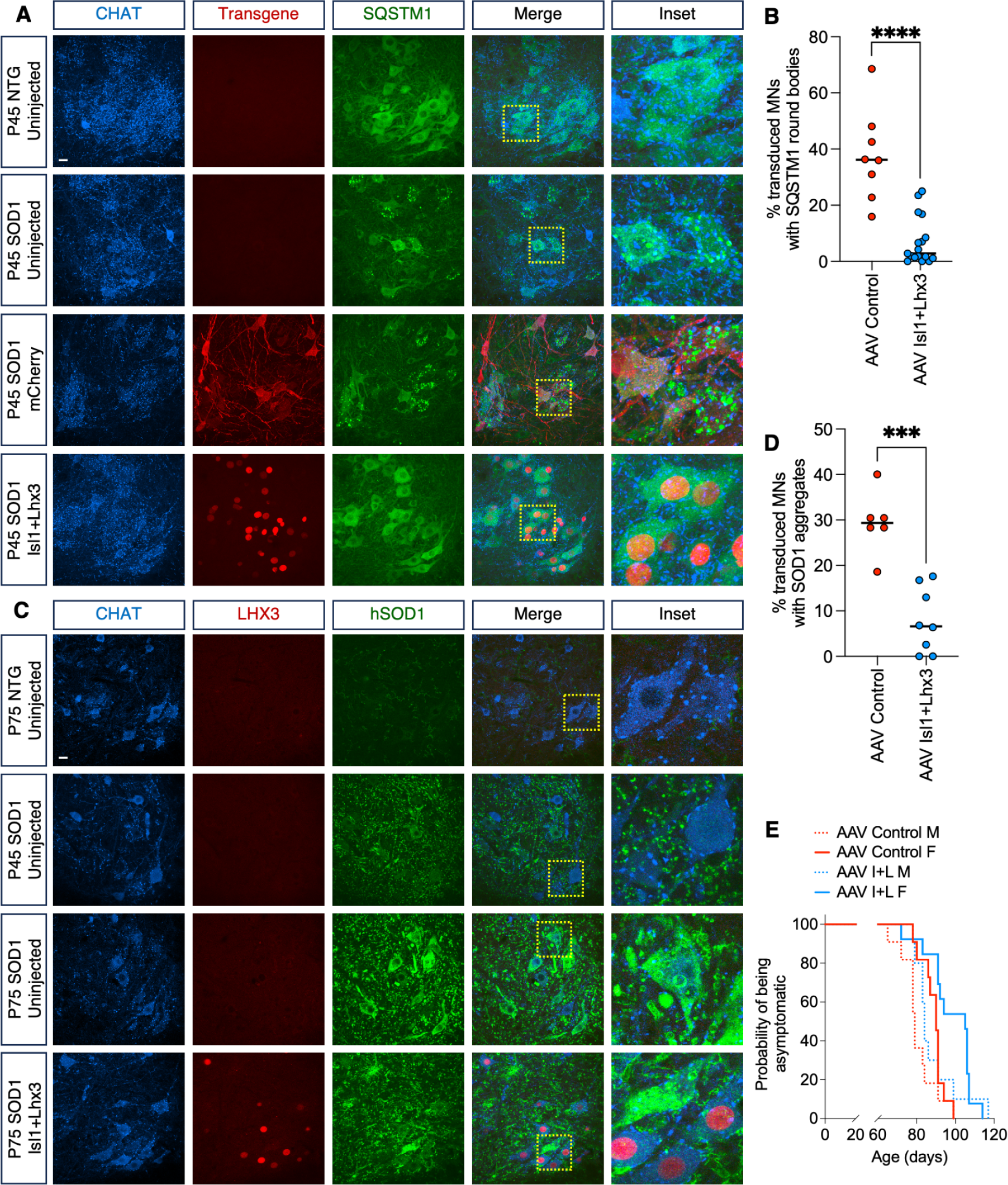
ISL1 and LHX3 re-expression attenuates ALS phenotypes in SOD1^G93A^ mice. (A) Representative immunostaining of SQSTM1 in L4-L5 ventral horn sections across treatment conditions in NTG and SOD1^G93A^ mice at P45. Transgene images represent immunostaining for LHX3 (NTG uninjected, SOD1 uninjected and SOD1 Isl1+Lhx3) or RFP (SOD1 mCherry). Regions within dashed yellow boxes are enlarged in Inset. Images are maximum intensity projections of 15 z sections taken 3 µ apart. Scale bar represents 10 µm. (B) Quantification of SQSTM1 round bodies in AAV-transduced motor neurons in the L4-L5 region of the spinal cord in AAV-mCherry or AAV-Isl1+AAV-Lhx3-treated SOD1^G93A^ animals. The percentage of CHAT+/mCherry+ or CHAT+/LHX3+ motor neurons exhibiting SQSTM1 round bodies was quantified in 15 confocal sections from 6 70µ hemisections per animal. Each point represents the mean percentage per hemisection from one animal. Black bars represent animal medians per treatment condition. Significance determined by unpaired two-tailed t test (p<0.0001; t=6.290, df=23). (C) Representative immunostaining of human SOD1 in L4-L5 ventral horn sections across treatment conditions and time. Regions within dashed yellow boxes are enlarged in Inset. Images are maximum intensity projections of 3 z sections taken 3 µ apart. Scale bar represents 10 µm. (D) Quantification of SOD1 aggregation in AAV-transduced motor neurons in the L4-L5 region of the spinal cord in AAV-mCherry or AAV-Isl1+AAV-Lhx3-treated SOD1^G93A^ animals at P75. The percentage of CHAT+/mCherry+ or CHAT+/LHX3+ motor neurons exhibiting SOD1 aggregates was quantified in 15 confocal sections from 6 70µ hemisections per animal. Each point represents the mean percentage per hemisection from one animal. Black bars represent animal medians per treatment condition. Significance determined by unpaired two-tailed t test (p=0.0001, t=5.669, df=12). (E) The probability of tremor onset in AAV-mCherry vs AAV-Isl1+AAV-Lhx3-treated SOD1^G93A^ mice was compared by Log-rank (Mantel-Cox) tests (n=10-13 animals per treatment per sex). Separate analyses were performed for male vs. female animals. For females, AAV-Isl1+AAV-Lhx3 treatment delayed tremor onset by 15 days (p=0.0036). For males, tremor onset was delayed by 5 days but was not significant (p=0.1138).

We evaluated the effects of ISL1 and LHX3 re-expression across a range of viral doses (6.37E+10 – 3.69E+11 vg/animal) on SQSTM1 round body formation in SOD1^G93A^ animals treated with AAV-Isl1+AAV-Lhx3 at P1 and analyzed at P45. Across all doses, we found that the percentage of AAV-transduced motor neurons exhibiting SQSTM1 round bodies was strongly reduced in AAV-Isl1+Lhx3-treated animals compared to mCherry (2.86% vs. 36.18%; p<0.0001, Fig. 4, A and B). We also found that the total number of motor neurons per hemisection that exhibited SQSTM1 pathology was dose-dependent and correlated significantly with motor neuron transduction efficiency (R^2^=0.5910, p=0.0003; Fig. S7A). At transduction efficiencies greater than ∼80%, SQSTM1 round bodies were almost completely abrogated (Fig. 4, A and B, and Fig. S7A), pointing to a cell-autonomous effect of ISL1 and LHX3 re-expression on SQSTM1 pathology. AAV-Isl1+AAV-Lhx3 treatment did not reduce the average number of motor neurons per hemisection (data not shown), nor did it prevent the expression of *Sqstm1* (Fig. 4A and Fig. S7B). Furthermore, the percentage of motor neurons exhibiting SQSTM1 pathology did not differ between AAV-mCherry-treated and uninjected controls (Fig. 4A), indicating that round body formation was not affected by AAV transduction itself, nor did the ectopic expression of mCherry exacerbate it. Taken together, these findings suggest that the effect of ISL1 and LHX3 re-expression on SQSTM1 round body formation in transduced motor neurons was a direct reflection of enhanced resilience to stress resulting from mutant SOD1 expression.

### ISL1 and LHX3 factors suppress the formation of pathological SOD1 protein aggregates in spinal motor neurons

SQSTM1 has been shown to selectively bind mutant SOD1 and actively sequester it into insoluble cytoplasmic inclusions of ubiquitinated proteins that intensify over time in spinal motor neurons in SOD1 mutant mouse models ^16,17^. To investigate the effects of ISL1 and LHX3 re-expression on this process, we quantified the incidence of SOD1-positive aggregates in spinal motor neurons using an antibody specific for the human transgene. Compared to SQSTM1 aggregates, which were pronounced by P45, we observed that cytoplasmic SOD1 aggregates were more readily detectable at later stages of the disease (Fig. 4C), consistent with previous reports ^20^. By P75, when SOD1 aggregation was severe in untreated or AAV-mCherry-treated animals, we found that AAV-Isl1+AAV-Lhx3 treatment reduced the number of cells positive for SOD1 aggregates by >3-fold in transduced motor neurons (p=0.0042; Fig. 4D). Given that AAV-Isl1+AAV-Lhx3 treatment did not alter the expression of endogenous *Sod1* in INTACT mice (Fig. S7C), it is unlikely that the effects of ISL1 and LHX3 re-expression on transgenic SOD1 aggregation are the result of reduced *Sod1* expression. Accordingly, mutant SOD1 can still be detected by IHC in AAV-Isl1+AAV-Lhx3-transduced motor neurons, albeit in a diffuse pattern that resembles uninjected or AAV-mCherry-treated animals at earlier stages of the disease (Fig. 4C). Together, these findings support the idea that ISL1 and LHX3 re-expression in mature motor neurons can prevent the cascade of proteostatic deficits that stems from the overexpression of mutant SOD1 and drives disease progression.

### ISL1 and LHX3 delay the onset of clinical disease in ALS mice

Considering that the exacerbation of SQSTM1-linked protein dyshomeostasis accelerates the onset of clinical symptoms in SOD1 mutant mice ^15,17^, we then asked whether AAV-Isl1+AAV-Lhx3 treatment could delay symptom onset. For these experiments we transduced mice at P1 with 7.56E+10 vg/animal, yielding ∼22% AAV+ motor neurons by P45 (Fig. 1D). SOD1^G93A^ animals injected with AAV-mCherry or AAV-Isl1+AAV-Lhx3 were monitored over time for the appearance of fine hindlimb tremors, which marks the onset of clinical symptoms in this model ^21^. Despite the relatively low percentage of motor neurons expressing the transgenes at late postnatal time points, we found that AAV-Isl1+AAV-Lhx3 treatment significantly delayed the appearance of tremors in females from P90 to P105 (p=0.0036; Fig. 4E). There was a similar trend in males, though it was not significant (median delay of 5 days from P79 to P84; p=0.1138; Fig. 4E), perhaps because male SOD1^G93A^ mice generally show more rapid progression of clinical phenotypes than females ^22^. Given the dose-dependency of the effects of AAV-Isl1+AAV-Lhx3 on SQSTM1 aggregation, these findings raise the possibility that increasing the degree of motor neuron transduction and improving the stability of transgene expression in future studies could yield additional and longer-lasting phenotypic benefits.

## Discussion

This study is a first step toward the development of highly targeted reprogramming approaches to rejuvenate susceptible neuronal types and suppress adult-onset neurodegenerative diseases. In contrast to methods that rejuvenate cells through the overexpression of general pluripotency factors (OCT4, SOX2, and KLF4), our approach combines native motor neuron factors with a motor neuron-specific expression system to achieve more targeted and biologically relevant reprogramming of the cells most vulnerable to disease. Heterochronic expression of ISL1 and LHX3 in mature neurons leads to changes in both chromatin accessibility and gene expression in alpha and type 3 motor neurons that are consistent with reversion to a younger cellular state. Functionally, we found that ISL1 and LHX3 re-expression at an early postnatal stage could prevent key histological and clinical phenotypes in a mouse model of ALS, an incurable, adult-onset motor neuron degenerative disease.

Previous work in corticospinal tract (CST) motor neurons supports the idea that reversion to an immature transcriptional state is protective and can enhance the regenerative capacity of neurons. Following spinal cord lesion, CST motor neurons reactivate a host of developmentally-regulated genes as part of an endogenous response to injury, and prolonging this immature transcriptional state through treatment with grafts of spinal neural progenitor cells can promote axon regrowth and restore limb function ^23^. We noted that many of the shared, upregulated alpha and type 3 DEGs populating GO terms associated with neurogenesis were also upregulated in regenerating CST motor neurons. These include genes involved in axon guidance and branching (*Unc5d*, *Dcc*, and *Fgf13*), as well as genes involved in synaptic plasticity (*Grik1* and *Rgs7*). While endogenous expression of these shared pro-regenerative genes normally peaks between embryonic and early postnatal stages of development, some are still expressed in adult spinal motor neurons and have been implicated in the maintenance of adult neural connectivity and protection from synaptic and axonal degeneration, such as *Raph1* and *Syt1* ^24^. Together, these observations raise the possibility that ISL1 and LHX3 are not only reverting spinal motor neurons to a less mature state, but are activating a gene expression program that is conducive to adult neuron regeneration.

While AAV-Isl1 and AAV-Lxh3 appeared to be expressed in all three major motor neuron subtypes, we were surprised to find that changes in gene expression and chromatin accessibility were limited to – and diverged between – alpha and type 3 motor neurons. The mechanisms underlying the subtype-selective effects of these motor neuron selector transcription factors are not immediately clear, but may be related to basal differences in genome accessibility that facilitate or prevent ISL1 and LHX3 binding. Alternatively, each motor neuron subtype may endogenously express other transcription factors or co-factors that modulate the activity of ectopic ISL1 and LHX3. Nevertheless, the preferential effects of re-expressed ISL1 and LHX3 on alpha over gamma motor neurons may be particularly important in the context of ALS, where alpha motor neurons are highly vulnerable to disease while gamma motor neurons are resilient ^25^. Little is known about the fate of type 3 motor neurons in ALS, but single nucleus RNAseq and ATACseq studies of motor neurons in vivo, including our own, may inform future strategies to selectively interrogate type 3 motor neurons in ALS.

To better understand the therapeutic potential of ISL1 and LHX3 re-expression, future studies will be required to evaluate its effects on late-stage ALS phenotypes, including motor dysfunction, paralysis, and early death. To date, our ability to perform long-term studies in SOD1^G93A^ mice has been limited by the rapid downregulation of ISL1 after P1 injection. Under similar low titer transduction conditions, ChatE-driven mCherry was strongly and continuously expressed in most motor neurons at all ages examined, while ISL1 and LHX3 expression dropped precipitously after ∼2 weeks. These findings suggest that the re-expressed transcription factors are either silenced or degraded in a selective manner. Increasing viral titers helped sustain LHX3 but not ISL1 expression, raising the question of whether ectopic ISL1 might be dispensable in the reactivation of immature gene expression programs and the attenuation of ALS phenotypes. Thus, evaluating the efficacy of high-titer AAV-Lhx3 on its own will be an important next step. A second, more therapeutically relevant approach would be to transduce motor neurons with AAV-Isl1 and AAV-Lhx3 (or AAV-Lhx3 alone) at later postnatal timepoints, including after symptom onset. This would provide insight into whether motor neuron-targeted interventions can be beneficial at late stages of the disease when motor neuron degeneration is underway and non-cell-autonomous factors such as neuroinflammation contribute to clinical progression ^26^.

In summary, our work demonstrates that heterochronic re-expression of embryonic selector transcription factors in postnatal animals can cell type-specifically increase the resilience of susceptible neurons and ameliorate cellular and behavioral pathologies in adult-onset neurodegenerative diseases.

## Materials and Methods

### Cloning pAAVs

ChatE, Isl1, and Lhx3 were Gibson cloned into an AAV2 backbone (pAAV). The pAAV backbone contained an AAV2 ITR, a cloning site, a mini promoter (TAGAGGGTATATAATGGAAGCTCGACTTCCAG), chimeric intron, mCherry, WPRE, SV40 poly(A) signal, and the second AAV2 ITR. This pAAV was linearized by digesting at the cloning site with Kpn1. ChatE was amplified from genomic DNA of wildtype C57BL6/J mice with primers containing ∼28 bps of homology to the pAAV backbone and inserted into linearized pAAV with Gibson cloning. This pAAV plasmid containing the ChatE and mCherry was then digested with BamHI and EcoRV to remove the mCherry and replace it with either Isl1 or Lhx3. Isl1 and Lhx3 cDNAs were amplified from other plasmids with ∼28 bps of homology to the pAAV-ChatE backbone for Gibson cloning. All digestion sites were recreated after Gibson insertions to make it easy to switch either the enhancer or the cargo in the pAAV plasmid.

The sequence of ChatE is as follows: CAGTGAGCTTCATTATCACCTAACAGCTTCAGAGTGGGTGGTGGGTTTTGGATGACAACCTTTCTTCTCATTTTATTCAGTGGCCACACCGTGGCCTTAGTCTGATAAACCAAAAACCTGCTCCATTATGAATCAGTGCTGTGGGGAGTGGGTAGAGAGTGTGAAGTTCTGGGGTGGGGGAGTCTGGAGAGAGGGTGGGAGCAGCCATTCTGCAGCAGTGCCTTCTTGGGGTCATGGGTCTGTAGGTGCTGCTGTGGAGGGAGAGATCAGCCTATTCTGGCTTCATTTCTGAGCTGCAAACTGCCTGGGTGTCTGGAGAAGCAGGTTGGCGTGGTGGTTAGCAGTGCGTGGGCGGGGTTGCCCGCTCTTGATTTATGATTTCTTTGTCTCTGTGGAAGCACTTAAGTGCAGGCTTTAGTTCCAATGACACTCAGGAGCCTCTGGATTCCAGCACTGGGGATGGGGGTGGGGTAGAACGTTCTCAGGCCTCACCAACCCCTCCCCTGTGTGCTGCCTTTGGGAGAGTCCCAAGGCTTCAGCATTACTTAATTAATTAGGCCTCTACTGCTACATAGGCTCAGATTCAAAAGAACAGAGTGGCCCACGTCAGCCATTCCCGGAAAAGTCTGATGGCTGGAAGCCAGAGGACTATGTGTCTGCCTTGCTGCCCTTGGCCAGCCCATCCTGAATGCCCAGACTCGGACAATGGAGTAGGTACAGAAGGGTAAAGACAGTGTCTTCTGTACCAGTAAGTGGGCCCTGATCTGCTCTCTACAGCTTCCAGAGAAAGGGCCTGGCCAATGAGCGGCCTTTTGAGTAGCAGATACCTCACATGCATTCTGATAGAAAGCCTGGCCCCAGATCACTGTGACTTTAGCCCTCAGGTTTCTTTTGCACTTCAATTCAATGACTTCTTGAGGTTCATTTCCCTCTCCAAGATTTGCCACAGACCAGTGGTTCTCAACCTGTGGGTCACACCTCCTTTGGGGAAATTGAATGA)

### AAV Production

AAVs were generated and titered by following the protocol in Challis et al ^27^. No meaningful changes were made to the protocol. We used the PHP.eB cap plasmid and pHelper plasmid in combination with one of the three pAAV plasmids (mCherry, Isl1, or Lhx3) to generate each virus independently. If the titer of the viral preparations fell below 2E+12 vg/mL, they were respun through an Amicon filtration device to reduce overall volume and increase the final concentration to the desired titer.

### Animals

All mouse experimental procedures were approved by the Columbia University Medical Center Institutional Animal Care and Use Committee.

*INTACT mice*: SnSeq studies were performed in INTACT mice that were generated and maintained through crosses of Chat-Cre mice (B6.129S-Chat*^tm1(cre)Lowl^/*MwarJ, Jax 031661) with Sun1-GFP nuclear reporter mice (B6;129-*Gt(ROSA)26Sor^tm5(CAG-Sun1/sfGFP)Nat^*/J; Jax 021039).

*SOD1^G93A^ mice*: All SOD1^G93A^ studies were performed in mice that were heterozygous for the mutant human SOD1^G93A^ transgene on a C57BL/6J background (B6.Cg-Tg(SOD1*G93A)1Gur/J; Jax 004435) and their non-transgenic littermates. Breeding pairs consisted of C57BL6/J females (Jax 000664) and SOD1^G93A^ males purchased from Jackson laboratories, where they were assessed for transgene copy number maintenance. Pups were genotyped at P0 and balanced into treatment groups by sex, genotype, and littermate status.

### Intracerebroventricular AAV injections

Intracerebroventricular AAV injections were performed on cryoanesthetized pups at P1 using a 10 µL syringe (Hamilton 7653-01) outfitted with a 0.375-inch 32-gauge needle (Hamilton 7803-04). The needle was inserted at 2/5ths of the distance between the lambda suture and the middle of the eye at a depth of 3 mm, and up to 6 µL of virus was injected per animal.

### Immunohistochemistry

For all immunohistochemical analyses, tissues were collected by transcardial perfusion with 10 mL ice-cold phosphate-buffered saline, followed by 40 mL of 4% paraformaldehyde in 0.1M phosphate buffer. The skull and spine were gross dissected and post-fixed for 18-24h in 4% paraformaldehyde. The L4-L5 region of the spinal cord was isolated by cutting the spinal cord at the T10 and S1 vertebrae, followed by laminectomy and transection of the spinal cord at the L3 and L6 roots. Spinal cord segments were stored at 4° in PBS containing 0.1% sodium azide until they were sectioned at 70 µ using a vibratome fitted with a disposable blade (Leica). Immunostaining was performed in floating sections using antibodies against CHAT (1:250, Millipore AB144P), ISL1 (1:10000, Thomas M. Jessell Laboratory, Columbia University), LHX3 (1:5000, Thomas M. Jessell Laboratory, Columbia University), MNX1 (1:15000, Thomas M. Jessell Laboratory, Columbia University), CADPS2 (1:1000, Thermo Fisher PA5-20518), SQSTM1 (1:500, Abcam ab56416), or human SOD1 (1:250, R&D Systems MAG 3418). Sections were incubated for 18-24h in primary antibody solution containing 1% bovine serum albumin, 0.4% Triton, and 0.1% sodium azide in TRIS-buffered saline. Sections were then incubated overnight in a secondary antibody solution containing 0.4% Triton in TRIS-buffered saline using antibodies raised in donkey and conjugated to Alexa Fluor 405, 488, 594, or 647 (Thermo Fisher). Sections were washed, mounted on slides, and coverslipped with Fluoromount-G (Southern Biotech).

### Confocal Microscopy

Immunostained spinal cord sections were visualized at 5x using an epifluorescent microscope (Zeiss Axioscope) and at 40x using a confocal microscope with an oil objective (Zeiss LSM 900). For confocal microscopy, z-sections were collected every 3µ.

### Image quantification

An experimenter blind to the treatment and genotype of the animals performed manual quantification of MNX1 frequency, SQSTM1 round body formation, and SOD1 aggregation on confocal images. Quantification of CADPS2 positive nuclei and CADPS2 intensity was performed using custom automated macros in ImageJ (source code available upon request). Statistical analyses were performed in Prism (v. 10).

### Bulk seq procedure + analysis

INTACT mice (n=4) were injected with AAV-Isl1+AAV-Lhx3 (6.38 total vg/animal) at P1. Sun1-GFP + nuclei were collected at P21 using a bead-based IP method described in detail in Patel et al. ^2^. Bulk RNA was isolated from FACs nuclei using trizol-based phase separation and the Zymo MicroPrep RNA kit. Libraries were prepared at the MIT MicoBio Center using SMARTer Stranded Total RNA-Seq Kit v2 - Pico Input Mammalian kit and sequenced paired end using the 150nt Nextseq kit. Reads were trimmed for adaptors and low-quality positions using Trimgalore (Cutadapt v0.6.2) ^28^. Reads were aligned to the mouse genome (mm10) and gene-level counts were quantified using RSEM (v1.3.0) ^29^ rsem-calculate-expression using default parameters and STAR (v2.5.2b) for alignment. Differential gene expression analysis was performed between injected animals and control P21 gene expression data ^2^ on RSEM gene-level read counts using EdgeR ^30,31^. For each set of comparisons being performed, genes were filtered for those that are expressed with CPM of ≥5 in all replicates in at least one condition.

### Single Nuc Seq procedure + analysis

Sun1-sfGFP-Myc mice were either injected with AAV-Isl1+AAV-Lhx3 at P1 (6 mice, Treatment group) or left untreated (8 mice, Control group). At P21 GFP+ nuclei were separately FAC sorted from treatment and control mice. Brachial and lumbar regions of the spinal cord were dissection and nuclei were isolated and sorted following the exact protocol detailed in Patel et al.^2^. All nuclei collected (∼60,000 from six treatment mice and ∼80,000 from eight control mice) were taken to the Genome Center at Columbia University, where they were counted, and processed using 10x multiome kit and 10x Chromium. Data was pre-processed using CellRanger by the Genomics Core. Alignment of sequencing reads were performed using both the standard mm10 genome and a custom genome that included 4 extra genes present in the AAV driven mRNA-the hIsl1 and hLhx3 coding sequences, chimeric-intron, and WPRE. Standard Seurat and Signac pipelines were then used to cluster and analyze the data. High-quality cells were first filtered using the following parameters: nCount_ATAC<7e4 & nCount_ATAC>5e3 & nCount_RNA<25000 & nCount_RNA>5000. Seurat objects were created from the remaining 7395 Control and 5108 Treatment cells. The two objects were integrated using canonical correlation analysis based on shared anchors identified in the snRNAseq data. 27 clusters were identified (dims = 1:20), each of which included both control and treatment nuclei. A previous study that performed snRNAseq on GFP+ nuclei from adult Chat-Cre; Sun1-sfGFP-Myc mice found clusters that consisted of Gad1 expressing GABAergic neurons, Slc17a6 expressing glutaminergic neurons, in addition to Chat expressing cholinergic neurons ^8^. Consistent with this previous study, we find a similar distribution of neurons in our clusters. Among the cholinergic clusters, we could identify three distinct motor neuron clusters, which correspond to alpha motor neurons, gamma motor neurons, and beta-like type 3 motor neurons based on markers identified in previous single nuclei RNAseq data ^8,9^. Cholinergic skeletal motor neurons (2094 cells) and all other cholinergic cells (1908) were then subsetted separately into new objects and reclustered without the use of canonical correlation-based integration (dims = 1:15 for both).

WPRE transcripts are used to show which clusters express AAV because Isl1 and Lhx3 transcripts could not be accurately quantified. Human Isl1 and Lhx3 genes were driven by ChatE-AAV. Sequencing reads that aligned only to the human cDNA were counted as AAV-driven transcripts and reads that align only to the mouse Isl1 and Lhx3 loci counted as endogenous; however any reads that aligned to both human and mouse genes were thrown out, making the quantification of these genes inaccurate.

All differential gene expression was performed using FindMarkers with a minimum log2FC of 0.25 and an adjusted p-value of 0.05. To analyze the ATAC data, peaks were called using Macs2 on clusters in the cholinergic skeletal motor neuron object. Differential peaks were identified using FindMarkers (test.use = ‘LR’, latent.vars = ‘nCount_peaks’). ClosestFeature was used to identify genes closest to peaks. RegionMatrix and RegionsHeatmap were used to make heatmaps of ATAC-seq reads in Fig. 3D. FindMotifs was used to perform motif enrichment on Jaspar mouse motif dataset (species = 10090).

GO analysis was performed for up-regulated alpha and type 3 DEGs using the STRING database v12.0 (string-db.org) against the background of all genes that were detected in the snSEQ dataset (9482 genes). FDRs <0.05 were considered significant.

### Tremor analysis in SOD1^G93A^ mice

For analysis of clinical onset, SOD1^G93A^ animals (n=10-13 per treatment per sex) were monitored for the appearance of hindlimb tremors on a daily basis beginning at P40. Tremors were assessed qualitatively as per Hatzipetros et al. ^21^ by an experimenter blind to the genotype and treatment of the animals. The probability of tremor onset in AAV-mCherry vs AAV-Isl1+AAV-Lhx3-treated SOD1^G93A^ mice was compared by Log-rank (Mantel-Cox) tests in Prism (v.10). Separate analyses were performed for male vs. female animals.

## Acknowledgements

The Virology Core at the Zuckerman Institute of Columbia University, led by David Ng, aided in viral production. Yael Weitzner was instrumental in validating the staining and quantification protocols for SQSTM1 round bodies. The illustrations in Fig. 1B were created in Biorender.

## Funding

This work was supported by funding from Project ALS and by NIH grants R01NS116141, R01NS109217, and K99NS121136.

## Author contributions

ERL, TP, and HW designed all experiments. ERL, TP, and HW wrote the manuscript and prepared the figures. TP designed and cloned the ChatE and AAV-Isl1 and AAV-Lhx3 constructs. GN, FL, EC, SA, and TP prepared the AAVs. ERL, EC, and GN injected pups with AAVs. EC, ST, and HM prepared tissue for immunohistochemistry. ST and ID performed immunohistochemistry and confocal imaging. EC, ST, and ID quantified HB9, CADPS2, SQSTM1, and SOD1 staining. HM assessed tremor onset. ERL performed all statistical analyses for histological and behavioral analyses. TP isolated nuclei for single nuc seq. TP, JAC, and ERL analyzed single nuc seq data.

## Competing interests

A U.S. Provisional Application has been filed for the use of ISL1/LHX3 in neurodegeneration (no. 63/359,527).

## Data and materials availability

All sequencing data in this study will be deposited to the GEO database.

---

**Fig. S1:**
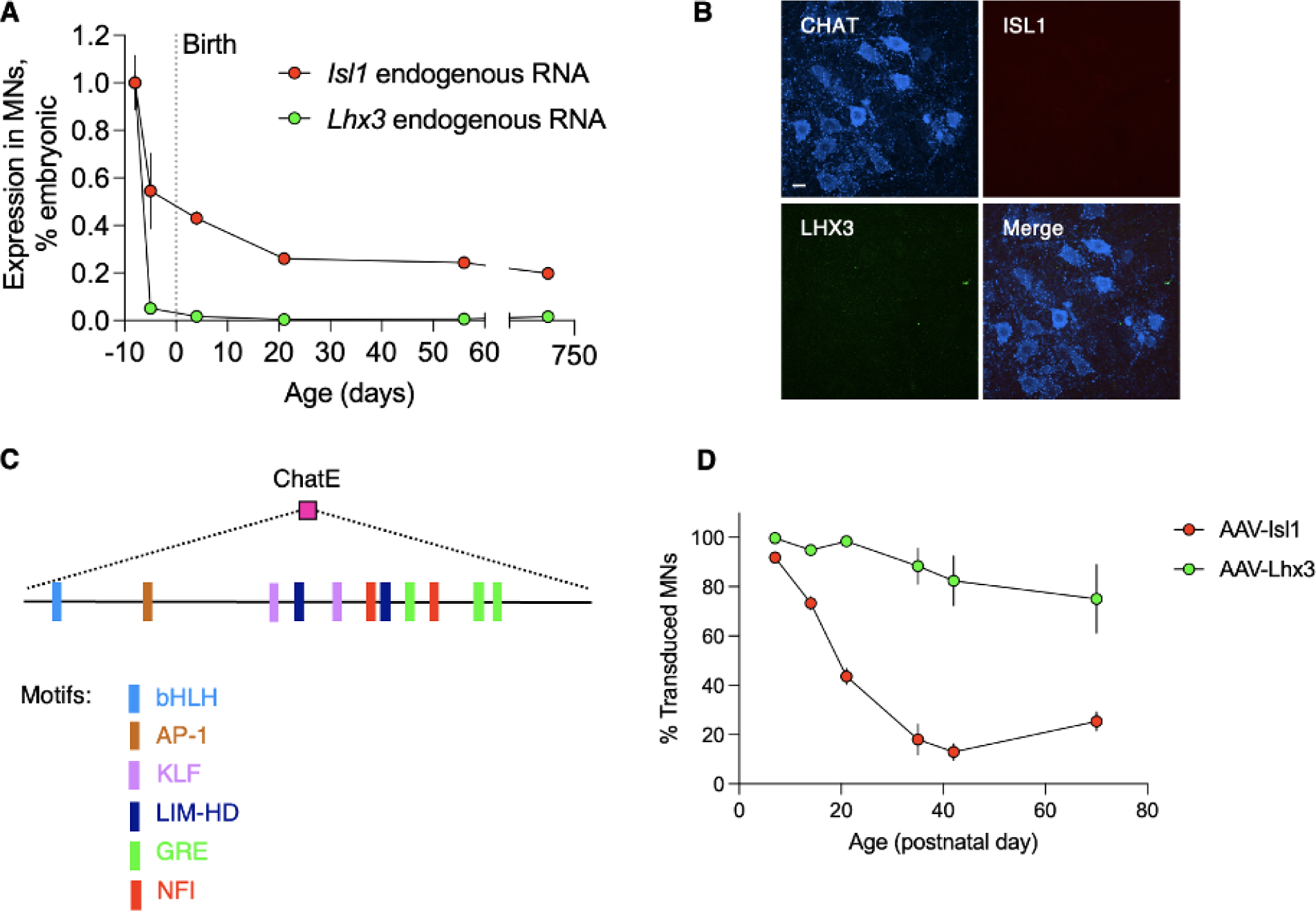
(**A**) Endogenous *Isl1* and *Lhx3* expression decrease sharply after their peak at E13.5 (RNAseq data from ^2^; expression for each time point normalized to values at E13.5). (**B**) Endogenous ISL1 and LHX3 are undetectable by immunofluorescence in CHAT+ cells in the L4-L5 ventral horn in untreated animals at P45. Images are maximum intensity projections of 15 z sections taken 3 µ apart. Scale bar represents 10 µM. (**C**) Schematic of the 1000 bp ChatE demonstrating the presence of transcription factor binding motifs enriched during motor neuron specification or maturation. (**D**) Quantification of transduced motor neurons in L4-L5 ventral horn at P45 in animals treated with AAV-Isl1 or AAV-Lhx3 (3.6E+11 - 3.8E+11 vg/animal) at P1. Sample size (n) was 2-4 animals per treatment per time point. Points represent mean transduced motor neurons per hemisection per animal, quantified across 6 70µ hemisections per animal from 15 confocal images per hemisection. Error bars represent SEM.

**Fig. S2:**
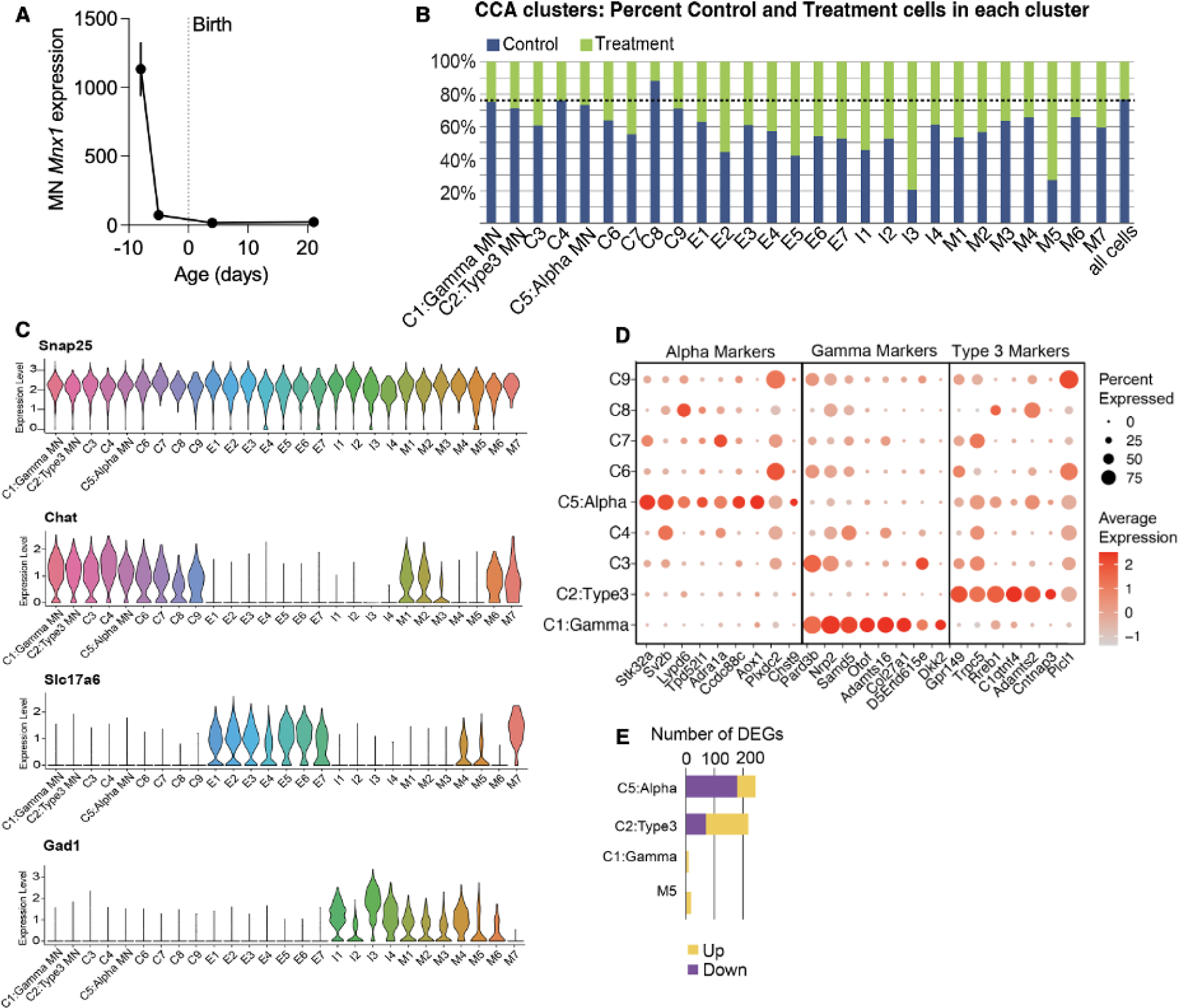
(**A**) RNAseq-derived expression levels of Mnx1 in mouse motor neurons over time ^2^. Numbers on the y-axis represent expression values normalized across all genes in all samples. (**B**) Percent control and treatment cells in all clusters. While there is some variation between clusters, all clusters are comprised of both control and treatment cells. (**C)** Violin plot showing **e**xpression of the neuronal gene *Snap25*, the cholinergic gene *Chat*, the excitatory gene *Slc17a6*, and the inhibitory gene *Gad1* in clusters from Fig. 2C. (**D**) Dot plot showing expression levels and percent cells expressing known alpha, gamma, and type 3 markers in cholinergic clusters. (**E**) Differential gene expression between AAV+ treatment cells vs. control cells in motor neurons and M5 clusters. Gene expression changes are largely restricted to alpha and type 3 motor neurons.

**Fig. S3:**
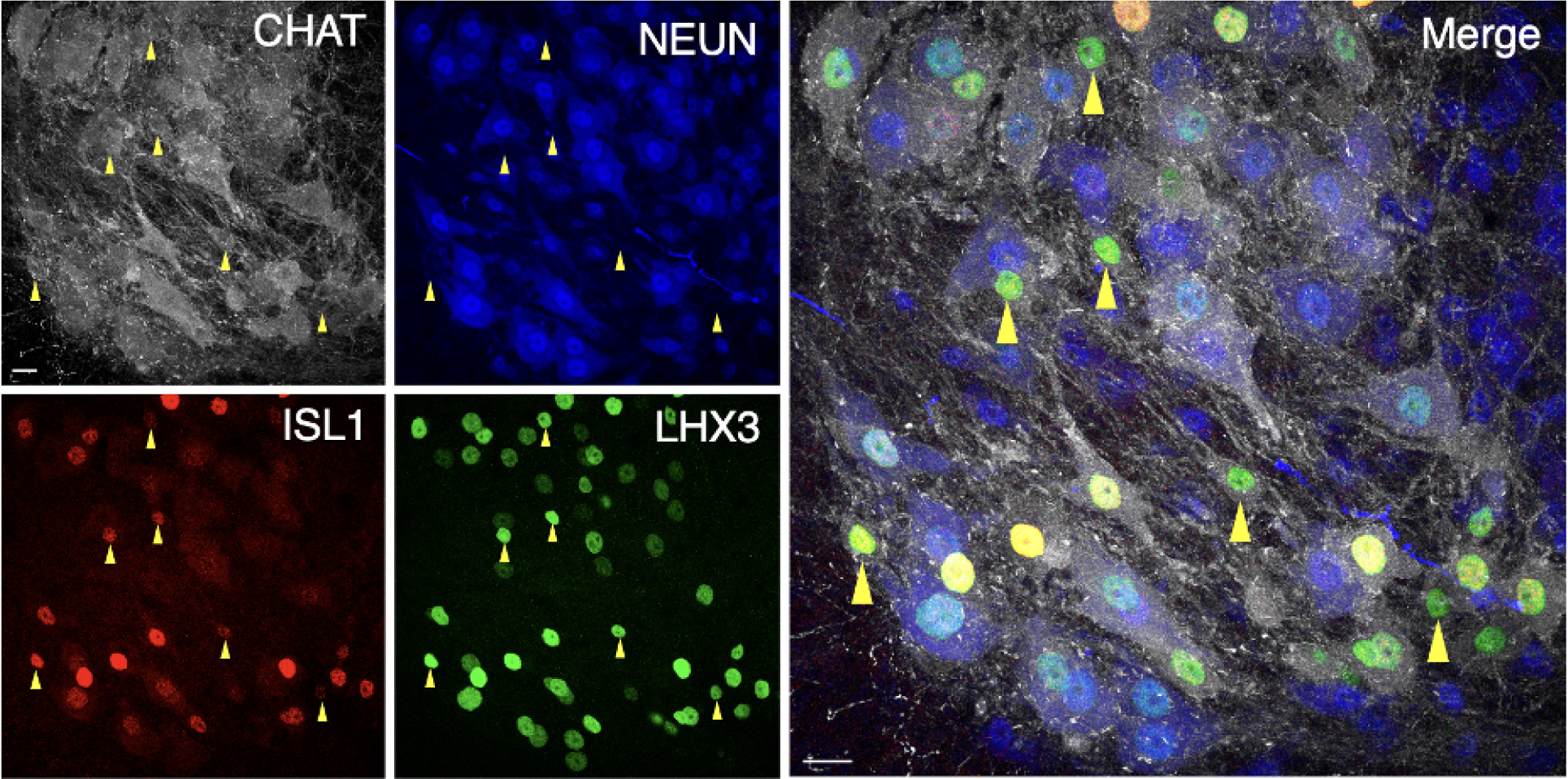
Representative immunostaining in the L4-L5 region of the ventral horn from animals injected with AAV-Isl1+AAV-Lhx3 (6.38E+10 total vg/animal) at P1 and analyzed at P14. Images are maximum intensity projections of 15 z sections taken 3 µ apart. Yellow arrowheads indicate AAV-transduced gamma motor neurons, identified as small-diameter CHAT+ cells that lack NEUN expression. Scale bars represent 10 µM.

**Fig. S4:**
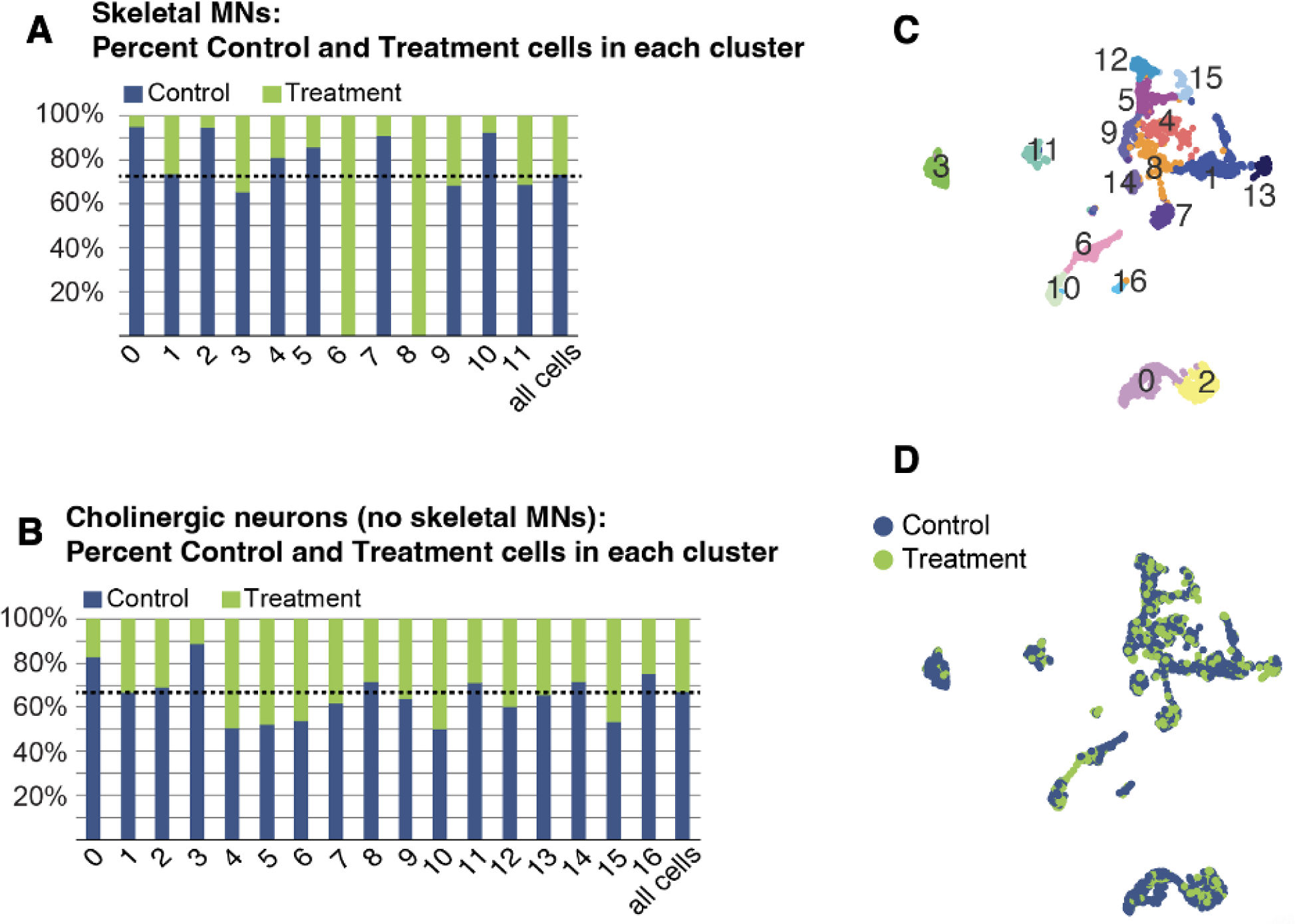
(**A**) Percent control and treatment cells in motor neuron-specific clusters. Two of the clusters, 6 and 8 (termed type 3 prime and alpha prime, respectively), are composed solely of treatment cells. These clusters represent type 3 and alpha motor neurons from the treatment group. (**B**) Percent control and treatment cells in non-motor neuron cholinergic clusters. While there is some variation between clusters, all clusters have both control and treatment cells. (**C-D**) Clustering of non-motor neuron cholinergic cells, and composition of control and treatment nuclei in each cluster.

**Fig. S5:**
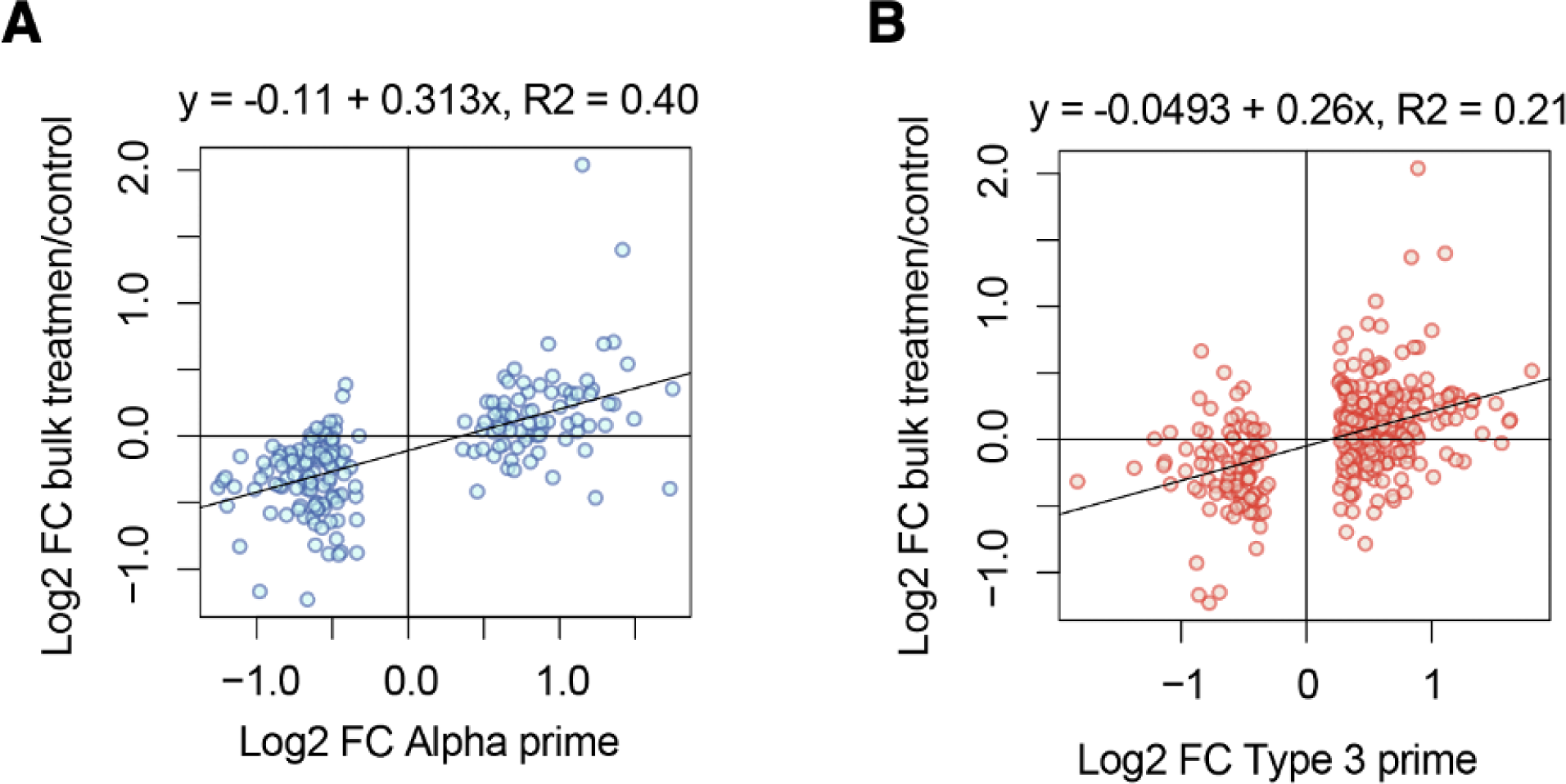
(**A)** Dotplot showing log2 fold expression change in alpha prime vs. alpha clusters (alpha DEGs) on x-axis and log2 fold expression change in the same genes in treated bulk RNAseq data vs. control bulk RNAseq data. There is an overall positive correlation in these gene expression changes despite differences in the methodology and sensitivity of snRNAseq vs. bulk RNAseq. (**B**) Same as (B) but with log2 fold expression change in type 3 prime vs. type 3 clusters (type 3 DEGs) on the x-axis. The positive correlation with bulk RNAseq is stronger for alpha than type 3 DEGs.

**Fig. S6:**
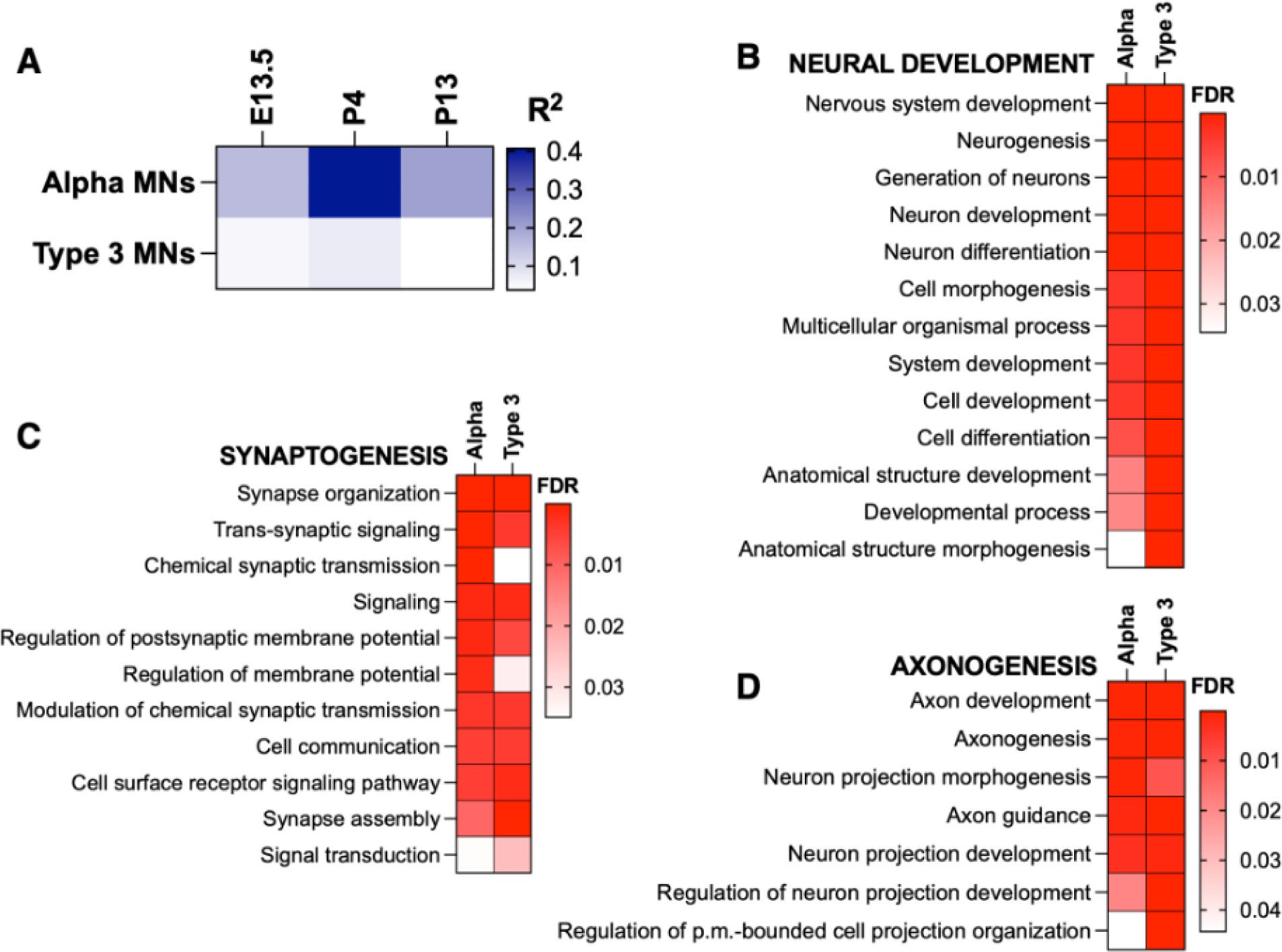
(**A**) Heatmap of R^2^ values for correlations between AAV-Isl1-Lhx3-mediated changes in gene expression vs. maturation-mediated changes in expression between P21 and the indicated time points for alpha and type 3 DEGs. Values for P4 are reported in Fig. 3G. P values for alpha DEGs are as follows: E13.5, p<0.0001; P4, p<0.0001; P13, p<0.0001. P values for type 3 DEGs are as follows: E13.5, p<0.0001; P4, p<0.0001; P13, p=0.0002. Minimum R^2^ value = 0.03823; maximum R^2^ value = 0.4063. (**B-D**) Heatmaps of FDR values for shared, significantly enriched GO terms between up-regulated alpha and type 3 DEGs relating to neural development (B), synaptogenesis (C), and axonogenesis (D). Maximum FDR value = 0.05; minimum FDR value = 1.29E-9.

**Fig. S7:**
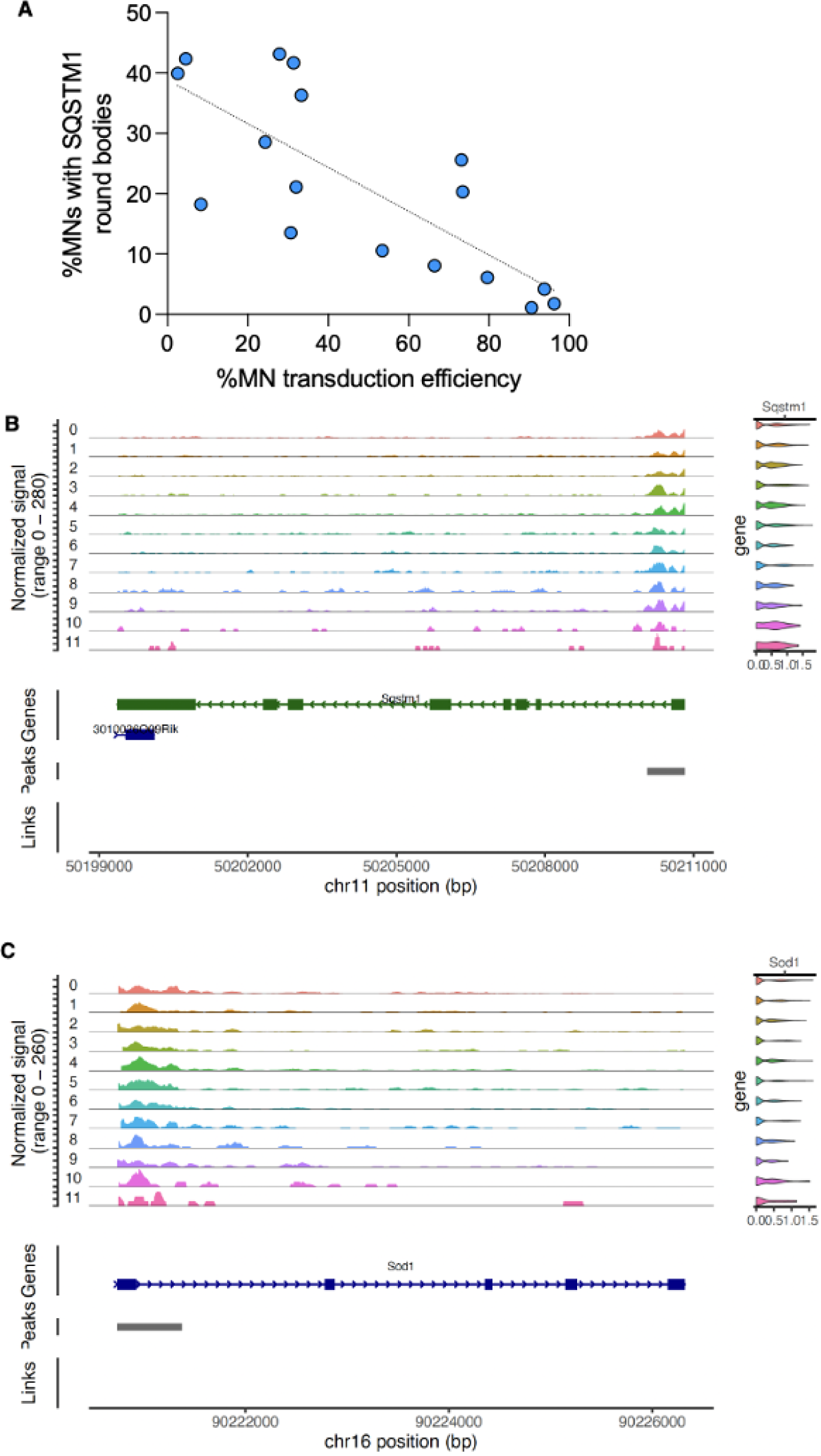
(**A**) Correlation between motor neuron transduction efficiency and incidence of SQSTM1 aggregates. Animals were treated with a range of AAV-Isl1+AAV-Lhx3 doses at P1 (6.37E+10 – 3.69E+11 vg/animal). Quantification of SQSTM1 round bodies and ISL1 and LHX3 expression in CHAT+ motor neurons at P45 was performed in 6 70µ L4-L5 hemisections per animal from 15 confocal images per hemisection. Each point represents one animal. Correlation between the percentage of all CHAT+ motor neurons per hemisection (transduced or not) exhibiting SQSTM1 round bodies was correlated with the percentage of CHAT+ motor neurons in the same hemisections that were transduced with AAV-Isl1 and/or AAV-Lhx3 by simple linear regression (R squared=0.5910; p=0.0003). (**B-C**) ATAC-seq reads at the genomic loci of Sqstm1 and Sod1 on the left, and violin plots showing gene expression on the right. Expression of these genes is not significantly different between treatment-specific clusters and control clusters.

